# *INTACT-olotl*: A simplified system for rapid cell type-specific profiling during axolotl limb regeneration

**DOI:** 10.64898/2026.09.18.752769

**Authors:** Jacob E. Barrett, Ramya Ailuri, Zinn Amos, Tim Curtis, Samantha Rock, Joshua D. Currie

## Abstract

Regenerating tissues like the axolotl (*Ambystoma mexicanum*) limb draw from many cell types, but isolating specific cell populations remains a bottleneck. Standard approaches rely on enzymatic dissociation and fluorescence-activated cell sorting (FACS), which are slow, costly, and require specialized instrumentation. We adapted INTACT (Isolation of <u>N</u>uclei <u>Ta</u>gged in specific <u>C</u>ell <u>T</u>ypes) to the axolotl, generating a transgenic line (*INTACT-olotl*) expressing *Sun1-sfGFP* in limb connective tissue, the population that carries limb positional information, dedifferentiates following injury, and forms the majority of the regenerative blastema. Combining an optimized nuclei isolation protocol with anti-GFP nanobody capture enriched labeled nuclei from ∼34.6% of input to 80–89% purity. Relative to dissociation and FACS, this recovers roughly 30-fold more target nuclei in less than half the time and without a cytometer. Bead-bound nuclei are directly applied to combinatorial-barcoding snRNA-seq, eliminating fluorescence-activated nuclei sorting (FANS). Profiling INTACT^+^ connective tissue during TGFβ inhibition revealed a loss of extracellular matrix remodeling with upregulated pro-fibrotic and inflammatory gene programs, resembling an unresolved wound environment. These changes reverse and regeneration resumes upon drug washout, indicating that functional TGFβ signaling is required for the transition from wound resolution to blastema formation.

## Introduction

Complex tissue regeneration requires the orchestrated effort of distinct cell types across days and weeks to resolve the initial inflammation, amass progenitors within a transient blastema, and coordinate the re-patterning of missing tissues (Sandoval-Guzmán and Currie, 2018). Although many heterogeneous cell types participate in regenerating structures like the axolotl salamander (*Ambystoma mexicanum*) limb, not all cell populations contribute in equal proportion or with equal importance to blastema formation (Currie et al., 2016). For instance, Lateral Plate Mesoderm-derived connective tissue makes up a large proportion of the limb blastema (estimated at 40-75%) and is thought to encode positional information in the patterned regenerate (García-García et al., 2025; Hu et al., 2026; Muneoka et al., 1986; Nacu et al., 2013). In contrast, muscle-derived cells represent a smaller subset of blastema cells and have little impact on successful regeneration (Hu et al., 2022; Sandoval-Guzmán et al., 2014).

Characterizing target cell types in vertebrate tissues typically involves enzymatic dissociation followed by fluorescence-activated cell sorting (FACS) of transgenic reporter-labeled cells (Gerber et al., 2018). However, this methodology is time intensive, requires access to specialized equipment, and can be cost-prohibitive for many labs. Both the process of enzymatic tissue digestion and time required for thorough dissociation and cell sorting contribute to known stress-related transcriptomic changes (Marsh et al., 2022; van den Brink et al., 2017). Unenriched cell suspensions are biased toward abundant, easily liberated cell types, so many more cells must be collected and sorted to recover rare or sought-after populations. Finally, the need for FACS instrument access can create financial and timing challenges for complex experiments.

An alternative strategy to isolate cell types of interest is through affinity isolation of transgene-labeled nuclei, referred to as INTACT *Isolation of <u>N</u>uclei <u>TA</u>gged in specific <u>C</u>ell <u>T</u>ypes*. The INTACT system was first implemented in *A. thaliana* through transgenic expression of bacterial *BirA* biotin ligase and the ligase recognition peptide, NTF, tethered to the nuclear envelope and driven by a cell type-specific promoter element (Deal and Henikoff, 2010). Subsequently, INTACT was adapted in Xenopus to characterize the developing heart proteome (Amin et al., 2014) and in mouse to affinity purify cell types in the heart, brain and liver (Bhattacharyya et al., 2019; Lawler et al., 2020; Loft et al., 2021). More recently deployed iterations of INTACT in vertebrates function by cell type-specific expression of the highly conserved nuclear envelope protein *Sun1* (*Sad1 and UNC84 domain-containing 1*) C-terminally fused to a fluorescent protein (FP). This allows for Sun1-FP-decorated nuclei to be purified using anti-FP antibodies for bulk genomics or transcriptomics (Mo et al., 2015). To achieve single nuclei transcriptomes, previous studies used INTACT-labeled nuclei as an input for FANS (Fluorescence-Activated Nuclei Sorting) without an affinity capture step, effectively equivalent to the lengthy cytometer sorting for dissociated cells (Alkaslasi et al., 2021; Su et al., 2024).

We adapted the INTACT system to axolotl by creating a Sun1-sfGFP transgenic that specifically labels limb connective tissue (INTACT-*olotl*). We generated an optimized nuclei isolation protocol and two nuclei precipitation workflows prioritizing either speed with INTACT enrichment or INTACT purity, slightly impacting time and yield. Benchmarked to standard tissue dissociation protocols, our INTACT-*olotl* protocol produces similar proportions of transgene-labeled nuclei but in a fraction of the time needed for enzymatic tissue digestion and with substantially improved yield. To demonstrate the utility of INTACT-*olotl*, we coupled INTACT nuclei enrichment with combinatorial barcoding snRNA-seq, creating an experimental workflow from tissue collection to single nuclei sequencing libraries that bypasses fluorescent cell sorting and can be performed in most molecular biology-equipped laboratories. We applied this methodology to profile connective tissue-centered transcriptional changes that occur at early stages of blastema formation following perturbation to TGFβ signaling. As the first application of INTACT in a highly regenerative species, we see this methodology as an accessible and flexible tool to efficiently characterize pro-regenerative cell types in the axolotl and newly emerging model organisms.

## Results

### Development of *INTACT-olotl* for profiling axolotl connective tissue cells during limb development, homeostasis, and regeneration

To adapt INTACT to axolotl and specifically label limb connective tissue cells, we first subcloned an existing INTACT cassette (Lawler et al., 2020) into a standard I-SceI and Tol2 transgenesis vector. The INTACT cassette contained an N-truncated mouse *Sun1* (*Sad1* and UNC84 domain containing 1) nuclear membrane protein (AA 208-757) which protrudes from the inner nuclear membrane into the perinuclear space (Fig. 1A-B). The *Sun1* C-terminus is fused to a copy of Superfolder GFP (sfGFP) (Fig. 1B). The INTACT cassette is driven by a well characterized 2.4Kb mouse *Prrx1* promoter/enhancer element (*mPrrx1enh::INTACT*) (Currie et al., 2019; Logan et al., 2002; Martin and Olson, 2000). The *mPrrx1enh* marks Lateral Plate Mesoderm-derived connective tissue (CT) cells in the axolotl limb during development, homeostasis, and regeneration (Gerber et al., 2018). Embryos injected with *mPrrx1enh::INTACT* and I-SceI meganuclease for integration were screened for fluorescence at limb bud stage 44/45 and axolotls possessing uniform expression in the mesenchyme were used for experiments or raised as F0 founders (Fig. 1C). We confirmed that Sun1-sfGFP localized to the nuclear membrane of CT nuclei with confocal microscopy (Fig. 1D-E) and that expression is maintained stably in mature connective tissue cells and not silenced in the 3-6cm (snout to cloaca) axolotls used for experiments (Fig. 1F). We performed distal amputations through the radius/ulna of axolotls and observed recruitment of *mPrrx1enh::INTACT^+^* cells to the blastema at 12 days post amputation (DPA) (Fig. 1G). Isolated nuclei from *mPrrx1enh::INTACT^+^* connective tissue cells retained sfGFP through tissue homogenization and purification (Fig. 1E).

**Figure 1.**
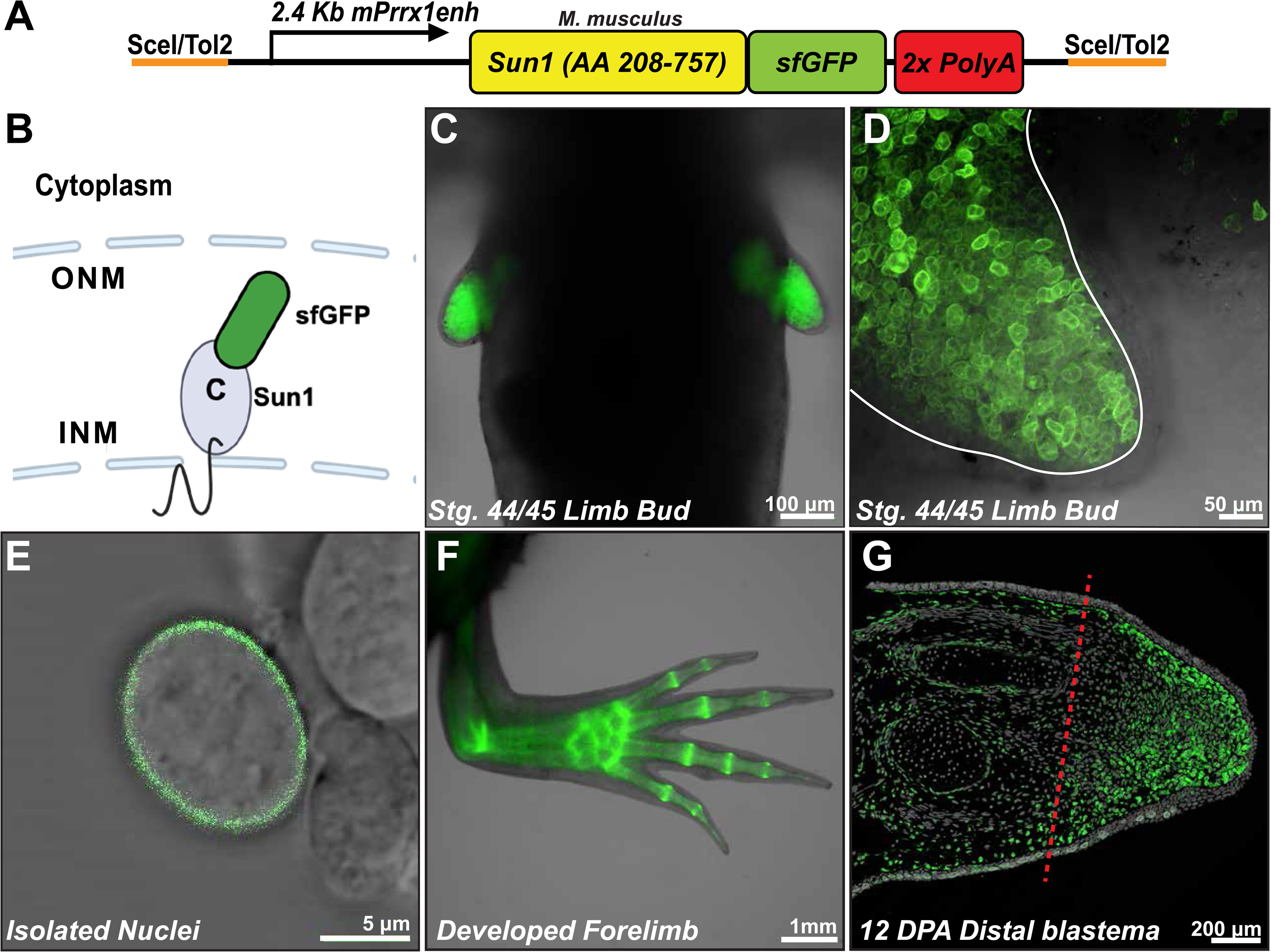
The *mPrrx1enh::INTACT* transgene labels the nuclear membrane of connective tissue (CT) nuclei during limb development, homeostasis, and regeneration. (**A**) Transgenic construct containing I-SceI/Tol2 integration sequences, the limb connective tissue-specific 2.4Kb mouse *Prrx1* promoter/enhancer element (*mPrrx1enh*), INTACT *Sun1* nuclear membrane protein (amino acids 208-757) fused to superfolder GFP (*sfGFP*). (**B**) Cartoon representation of Sun1-sfGFP fusion protein embedded in the inner nuclear membrane with sfGFP protruding into the perinuclear space (Created in BioRender. Currie, J. (2026) https://BioRender.com/lvxfo9e). (**C**) The *mPrrx1enh::INTACT* transgene labels Lateral Plate Mesoderm-derived progenitors during axolotl forelimb development (embryonic stage 44/45). (**D**) Higher magnification Stg. 44/45 limb buds with nuclear membrane sfGFP expression in limb bud mesenchyme. White line denotes epithelial-mesenchymal boundary. (**E**) Confocal imaging of isolated nuclei positive for Sun1-sfGFP following tissue homogenization and nuclei purification. (**F**) Developed forelimbs of 3-6 cm animals with connective tissue-specific *mPrrx1enh::INTACT* expression. (**G**) *mPrrx1enh::INTACT* lower limb blastema at 12 days post amputation. Red dashed line denotes the amputation plane.

### An optimized protocol yields high quality INTACT-labeled nuclei

The first step in the INTACT*-olotl* workflow consists of harvesting transgenic tissue followed by brief bead-based homogenization. Crude nuclei isolations are then filtered, pelleted, and washed. Resuspended nuclei are immunoprecipitated using anti-GFP nanobodies and can be used directly as input for downstream assays (Fig. 2A). A primary advantage of nuclei-based methodologies such as INTACT-*olotl* is that tissue homogenization and hypotonic lysis efficiently liberate nuclei, bypassing expensive, lengthy, and incomplete enzymatic digestion. To identify conditions which yield high-quality nuclei preps, we tested several previously published nuclei isolation protocols (Loft et al., 2021; Nadelmann et al., 2021; Wiegleb et al., 2022). We first processed tissue with a Dounce homogenizer as previously described (Loft et al., 2021; Wiegleb et al., 2022) and found that this method yielded preps with substantial debris and nuclei aggregation visualized with intense smears of trypan blue staining (Fig. 2B). Addition of 1mM DTT, 0.5mM spermidine, and 1x protease inhibitor immediately before Dounce homogenization produced a slight decrease in debris and nuclei aggregation (Fig. 2C). To further improve the process, we adapted a PBS and sucrose-based buffer system (Martin et al., 2023) with addition of DTT, spermidine, BSA, protease inhibitor, Igepal, and RNase inhibitor in axolotl tissue. This buffer, combined with a low-intensity bead-based homogenization step (Nadelmann et al., 2021) produced an optimized protocol which yielded high-quality preps with little to no debris or nuclei aggregation present (Fig. 2D-E, Supplemental Material). We observed that after isolation the nuclear membrane was fully intact and lacked membrane blebs that would indicate compromised membranes or suboptimal buffer osmolarity (Fig. 2D). Compared to established methods (Loft et al., 2021; Nadelman et al., 2021; Weigle et al., 2022) our optimized protocol yields the highest quality nuclei suspensions from axolotl tissue (Fig. 2E).

**Figure 2.**
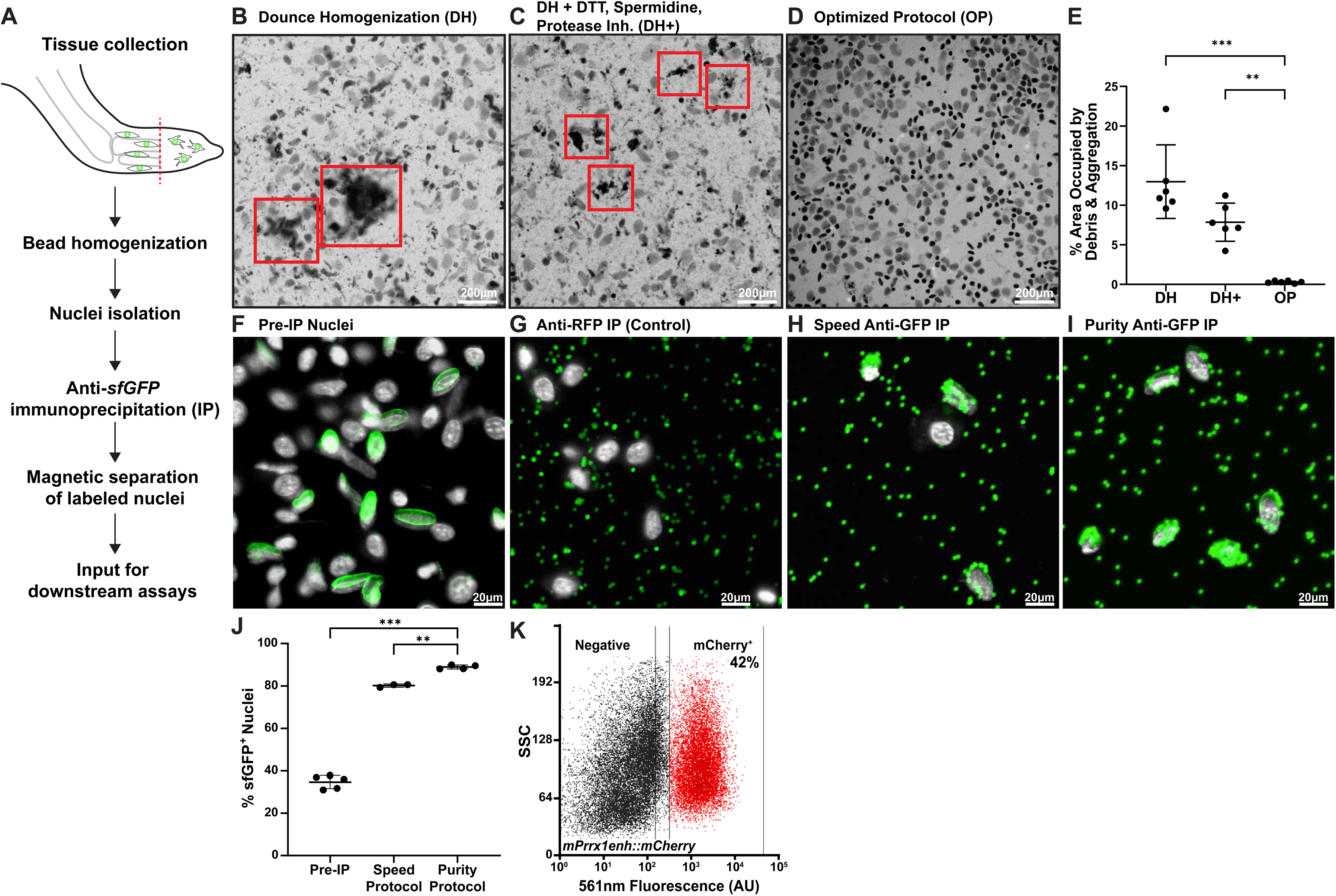
An optimized nuclei isolation protocol produces high quality, single nuclei suspensions that are efficiently enriched using anti-GFP conjugated nanobodies. (**A**) The INTACT-*olotl* nuclei isolation and purification workflow. (**B-E**) Tissue homogenization and nuclei isolation using (**B**) dounce homogenization alone (DH), (**C**) dounce homogenization plus the addition of DTT, spermidine, and protease inhibitor (DH +), or (**D**) an optimized protocol (OP) using bead-based homogenization and hypotonic lysis buffer. Trypan blue staining labels nuclei. Red boxes highlight areas of significant debris and nuclei aggregation. (**E**) Image aggregate quantification of trypan blue-stained nuclei isolation (n=6, *** = P < 0.001, ** = P < 0.002, One-way ANOVA, Tukey post hoc, data are mean +/SD). (**F**) Nuclei suspensions from *mPrrx1enh::INTACT* tissue yield heterogeneous populations of labeled (*Sun1-sfGFP*) and unlabeled (DAPI) nuclei before immunoprecipitation (IP). (**G**) Mock immunoprecipitation with anti-RFP nanobodies showing no binding of 488nm auto-fluorescent M-270 magnetic particles to DAPI-stained *mPrrx1enh::INTACT* nuclei. (**H-I**) Speed-IP and Purity-IP protocols enrich for *mPrrx1enh::INTACT* positive nuclei, visualized by anti-GFP auto-fluorescent M-270 magnetic particles covering Sun1-sfGFP nuclei periphery. (**J**) 34.6% (+/2%) of nuclei from *mPrrx1enh::INTACT* whole limb tissue are labeled prior to immunoprecipitation, Speed-IP and Purity-IP protocols yield final suspensions as quantified by confocal imaging of 80.2% (+/1%) and 88.9% (+/2%) *mPrrx1enh::INTACT^+^* nuclei, respectively, (N =3-5, n ≥ 75 nuclei, *** = P < 0.001, ** = P < 0.002, One-way ANOVA, Tukey post hoc, data are mean +/− SD). (**K**) Representative FACS plot of dissociated *mPrrx1enh::mCherry* axolotl limbs, with gated mCherry-positive cells making up 42% of total cells analyzed from the limb, comparable to the % of nuclei *mPrrx1enh::INTACT* positive from the limb.

We next established a workflow to rapidly enrich for nuclei expressing the INTACT transgene using anti-GFP nanobodies conjugated to a magnetic particle (ChromoTek). Prior to the purification, we confirmed that nuclei maintained sfGFP signal and normal nuclear morphology (Fig. 2F). To validate the specificity of the anti-GFP nanobody we performed a mock IP step where isolated INTACT nuclei were incubated with an anti-RFP nanobody. Visualization post-IP showed no enrichment of INTACT^+^ nuclei and no detectable binding to unlabeled nuclei (Fig. 2G). Based on iterative testing of precipitation conditions, we created two immunoprecipitation protocols: a rapid enrichment protocol (Speed-IP, Fig. 2H) and a high stringency, high purity protocol (Purity-IP, Fig. 2I). The protocols use the same homogenization steps and differ in the dilution of nuclei into multiple tubes and the number of post-IP wash steps. The Speed-IP protocol allows for nuclei to be enriched rapidly in a single tube and is designed to rapidly process large numbers of samples. The high stringency Purity-IP protocol dilutes sample nuclei across multiple tubes and increases the number of post-IP washes, conferring higher INTACT sample purity at the cost of increased post-IP time and a slight loss in yield. To quantify the efficiency of INTACT precipitation, we collected nuclei before and after the immunoprecipitation and quantified the percent of GFP^+^ nuclei present via confocal microscopy. Before purification approximately 34.6% of total nuclei were GFP^+^, with an enrichment to 80.2% and 88.9% of total nuclei being GFP^+^ for Speed-IP and Purity-IP protocols, respectively (Fig. 2F-J).

Current methods for isolating axolotl *mPrrx1enh^+^* limb cells rely on physical and enzymatic digestion of tissue followed by FACS (Gerber et al., 2018; Kawaguchi et al., 2024). To compare the INTACT-*olotl* workflow to previous FACS approaches, we quantified the percent enrichment between *mPrrx1enh::mCherry* enzymatic digestion and FACS-sorted cells and *mPrrx1enh::INTACT* nuclei isolation and precipitation. Dissociation and FACS of *mPrrx1enh::mCherry* limbs yielded 53,784 cells, 42% of which were mCherry^+^ (roughly, 22,500 cells, n = 6 limbs) (Fig. 2K, Table 1). Isolation of nuclei from equivalent-sized limbs yielded ∼1.8 million nuclei with ∼622,800 (34.6%, n = 6 limbs) being sfGFP^+^prior to immunoprecipitation. The INTACT-*olotl* workflow provides a 2.5-fold reduction in total experiment time per sample (FACS: 2.5 hours, INTACT-*olotl*: 1 hour) and a roughly 30-fold increase in target cell yield without cytometer sorting (Table 1).

**Table 1.**
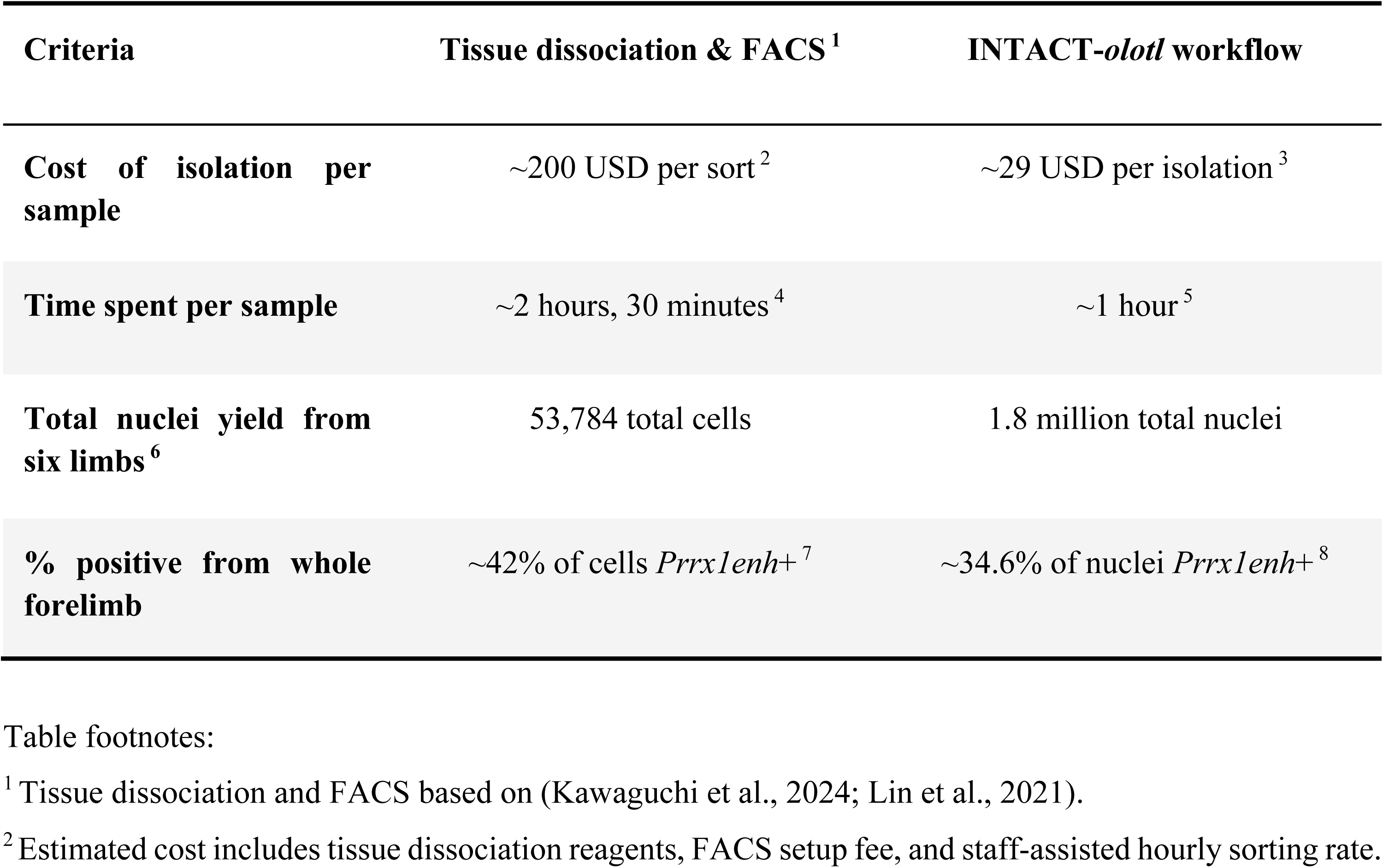
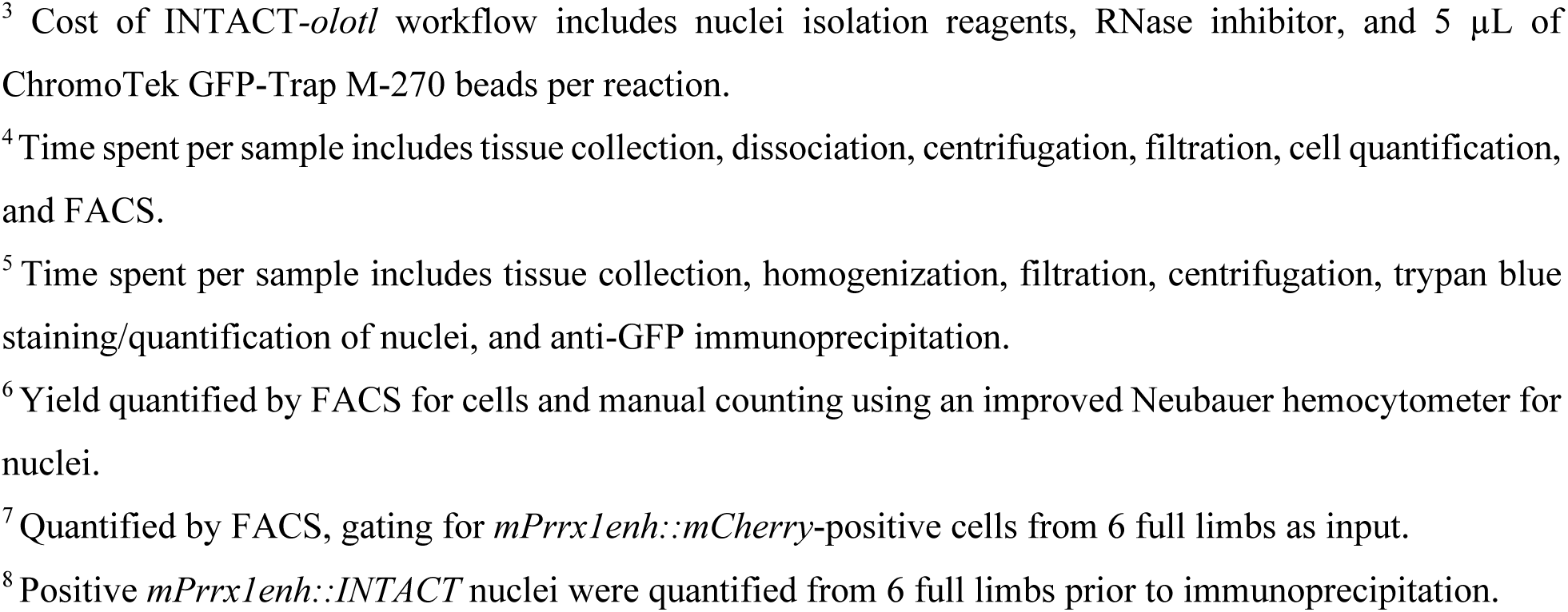
The INTACT-*olotl* system is more efficient, cost effective, and faster than standard FACS-based workflows.

| Criteria | Tissue dissociation & FACS<sup>1</sup> | INTACT-*olotl* workflow |
| --- | --- | --- |
| Cost of isolation per sample | ~200 USD per sort<sup>2</sup> | ~29 USD per isolation<sup>3</sup> |
| Time spent per sample | ~2 hours, 30 minutes<sup>4</sup> | ~1 hour<sup>5</sup> |
| Total nuclei yield from six limbs<sup>6</sup> | 53,784 total cells | 1.8 million total nuclei |
| % positive from whole forelimb | ~42% of cells *Prrx1enh*<sup>+</sup><sup>7</sup> | ~34.6% of nuclei *Prrx1enh*<sup>+</sup><sup>8</sup> |

Table footnotes:
<sup>1</sup> Tissue dissociation and FACS based on (Kawaguchi et al., 2024; Lin et al., 2021).
<sup>2</sup> Estimated cost includes tissue dissociation reagents, FACS setup fee, and staff-assisted hourly sorting rate.
<sup>3</sup> Cost of INTACT-*olotl* workflow includes nuclei isolation reagents, RNase inhibitor, and 5 μL of ChromoTek GFP-Trap M-270 beads per reaction.
<sup>4</sup> Time spent per sample includes tissue collection, dissociation, centrifugation, filtration, cell quantification, and FACS.
<sup>5</sup> Time spent per sample includes tissue collection, homogenization, filtration, centrifugation, trypan blue staining/quantification of nuclei, and anti-GFP immunoprecipitation.
<sup>6</sup> Yield quantified by FACS for cells and manual counting using an improved Neubauer hemocytometer for nuclei.
<sup>7</sup> Quantified by FACS, gating for *mPrrx1enh::mCherry*-positive cells from 6 full limbs as input.
<sup>8</sup> Positive *mPrrx1enh::INTACT* nuclei were quantified from 6 full limbs prior to immunoprecipitation.

### Single nuclei RNA sequencing of INTACT-*olotl* connective tissue in response to TGFβ inhibition reveals distinct transcriptional dynamics

To demonstrate the utility of the INTACT-*olotl* system, we performed single nuclei RNA sequencing (snRNA-seq) of mPrrx*1enh::INTACT^+^*-enriched nuclei across normal limb regeneration and Transforming Growth Factor Beta (TGFβ) pharmacological inhibition. TGFβ signaling is a conserved post-injury signal across both fibrotic and regenerative injury contexts (Finnson et al., 2013; Gilbert et al., 2013; Lakos et al., 2004; Lévesque et al., 2007). TGFβ is essential for limb regeneration (Lévesque et al., 2007); however high levels of TGFβ are correlated with fibrosis and sub-optimal healing outcomes (Deng et al., 2024), which is thought to be mediated through activation of inflammatory cytokines and inducing wound-activated fibroblasts to differentiate toward contractile myofibroblasts (Evans et al., 2003). The cognate axolotl limb cells, *mPrrx1enh*^+^ connective tissue, are thought to undergo a limited dedifferentiation post-injury toward a progenitor state (Lin et al., 2021). How TGFβ signaling functions during regeneration to transition post-injury inflammation toward blastema formation is unclear. Utilizing the INTACT-*olotl* system we sought to characterize pro-regenerative transcriptional dynamics downstream of TGFβ signaling.

We treated hemizygous F_1_ *mPrrx1enh::INTACT* axolotls daily with 25µM TGFβ inhibitor SB-431542 from 0 to 8 days post amputation (DPA) (Fig. 3A), which temporarily blocked blastema formation (Fig. 3B-C). Drug removal restored growth and blastema formation, albeit lagging morphologically compared to DMSO controls (Fig. 3B-C). Regenerating tissue (including 1 mm behind the amputation plane) was harvested at 0, 5, and 10 DPA (2 days after drug washout) for nuclei isolation and partial *mPrrx1enh::INTACT^+^* enrichment followed by processing for Parse Biosciences Evercode v3.0 snRNA-seq (Fig. S2). We identified 12 putative cell clusters encompassing connective-tissue, endothelial, epithelial, cycling, and immune cells (Fig. 3D). Although *Sun1-sfGFP* mRNA was detected in connective tissue clusters (Fig. S2), the overall penetrance across clusters was weak, consistent with prior scRNA-seq transgene detection, even when transgenes are driven from a strong, ubiquitous promoter (Gerber et al., 2018).

**Figure 3.**
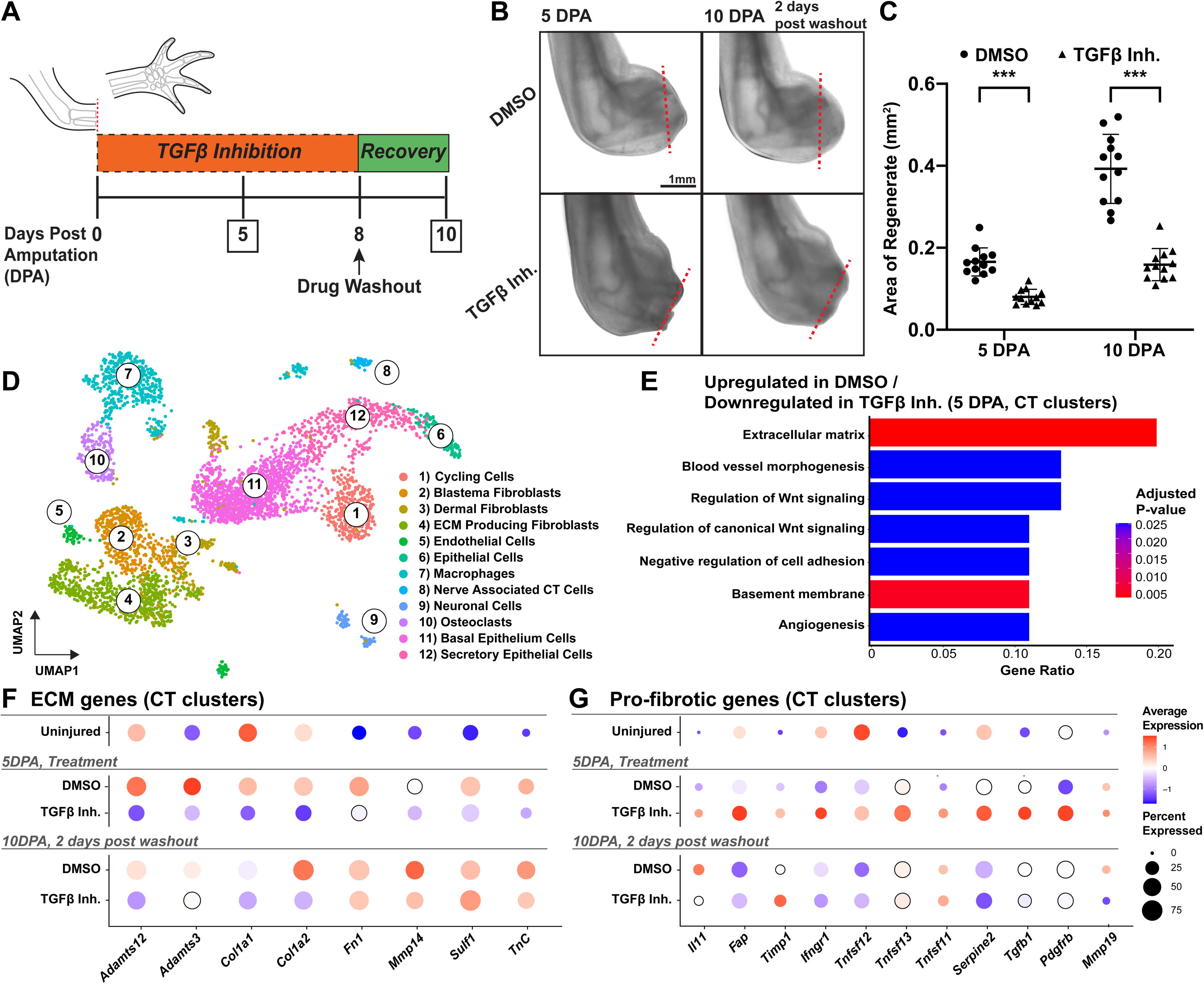
snRNA-seq of *mPrrx1enh::INTACT* enriched nuclei reveals significant changes to connective tissue cells in response to TGFβ inhibition. (**A**) Axolotls received either TGFβ inhibitor SB-431542 or DMSO for eight consecutive days following amputation. Tissue was collected at 0, 5, and 10 DPA, 2 days after drug washout (n = 4 per timepoint/treatment). (**B-C**) Representative images of regenerating axolotl limbs at 5 and 10 DPA following distal amputation. Red dashed lines represent the amputation plane. The area of regenerated tissue is significantly reduced under TGFβ inhibition at 5 and 10 DPA (n = 12, two-tailed Student’s t-test, *** = P < 0.001, data are mean +/SD). (**D**) Total UMAP across all timepoints (0, 5, 10 DPA) and treatments (DMSO, TGFβ-inhibited) for 4,432 nuclei identifies 12 distinct clusters including connective tissue, endothelial, epithelial, and immune cells. (**E**) GO analysis of CT-clusters at 5 DPA identifies processes upregulated in DMSO treated nuclei relative to TGFβ-inhibited nuclei. Color corresponds to adjusted p-values and gene ratio denotes the percentage of DEGs associated with a given GO term. (**F-G**) Dot plots depicting changes in the average expression of ECM and pro-fibrotic related genes across all timepoints (0, 5, 10 DPA) and conditions (DMSO, TGFβ Inh.) within the CT-clusters. Black outline is used to enhance visualization of dots with average expression around 0.

Within the connective tissue clusters, (Clusters 2, 3, 4, and 8, hereafter referred to as CT-clusters), TGFβ pathway components *Tgfbi*, *Ltbp3*, *Mfap2*, and *Smurf2* were downregulated, confirming that we had achieved effective pathway inhibition (Fig. S2). Gene Ontology (GO) analysis of the TGFβ-inhibited CT-clusters’ differentially expressed genes at 5 DPA highlighted downregulation of genes related to ECM formation, blood vessel morphogenesis, Wnt signaling, and regulation of cell adhesion (Fig. 3E). We identified extensive downregulation of both ECM structural components (*Col1a1*, *Col1a2*, *Fn1*, *Tnc*) and modifying enzymes (*Adamts12*, *Adamts3*, *Mmp14*, *Sulf1*) at 5 DPA in TGFβ inhibitor-treated CT-clusters (Fig. 3F). By 10 DPA and 2 days outside of drug treatment, TGFβ-inhibited CT-clusters had regained expression of many mis-regulated genes but maintained low expression of *Adamts12*, *Col1a1*, and *Col1a2* (Fig. 3F).

Surprisingly, we also observed an upregulation of typically pro-fibrotic genes in TGFβ-inhibited CT-clusters at 5 DPA. TGFβ-inhibited CT-cells exhibited high expression of *Fap* (Fibroblast activation protein), a marker of activated myofibroblasts correlated with fibrotic wound outcomes (He et al., 2024; Tillmanns et al., 2015), as well as upregulation of *Il11* (Interleukin 11), *Timp1, Ifng, Tnfsf11, Tnfsf12, Tnfsf13*, *Serpine2*, *Tgfb1*, *Pdgfrb*, and *Mmp19* (Fig. 3G). Expression of this pro-fibrotic gene program was largely resolved 2 days after drug washout at 10 DPA, suggesting that resumed TGFβ signaling was necessary to resolve injury-induced fibrotic gene expression and facilitate growth and blastema formation (Fig. 3G).

## Discussion

Single-cell and cell type-specific transcriptomics have provided increasing insight into the heterogeneity of complex tissues and morphogenetic processes. These techniques are becoming everyday tools in regenerative and developmental biology. However, the predominant workflows from living tissue to transcriptomic or genomic data require significant investment of time and reagents, complex experimental design, and access to specialized equipment such as cytometers. The alternative approach – unbiased cell isolation and sequencing – can dilute cell types of interest and lead to over-sampling of unwanted cell types. Previous scRNA-seq characterizations of axolotl regeneration have been complicated by over-representation of erythrocytes (Rodgers et al., 2020) and of epithelial cells, the latter requiring a selective removal of epithelia from harvested tissue (Leigh et al., 2018). Using an INTACT approach, we can alleviate a number of issues that hamper cell type-specific transcriptomics while optimizing time and cost. Our nuclei isolation and precipitation protocols let users prioritize either speed or purity. The Speed-IP option deliberately retains a fraction of unlabeled nuclei, which is useful when neighboring cell types are informative for snRNA-seq. Both protocols produce nuclei yields that outperform existing cell dissociation protocols (Fig. 2).

Our workflow is the first description of precipitated INTACT nuclei being directly used for single nuclei RNA-seq library preparation, completely eliminating fluorescence-activated nuclei sorting (FANS), a feature of previous INTACT workflows (Alkaslasi et al., 2021; Su et al., 2024). What makes this possible is the use of split-pool combinatorial barcoding strategies for snRNA-seq (e.g., SPLiT-seq, (Rosenberg et al., 2018)), where populations of nuclei are passed through serial barcoding and nuclei act as barcoding vessels while still bound to anti-GFP beads. This allows us to bypass microfluidic droplet-based barcoding. A further advantage is the minimal equipment burden. Apart from a bead homogenizer for tissue lysis, the entire procedure from tissue collection to snRNA-seq library preparation was performed using standard molecular biology bench equipment. The INTACT-*olotl* cassette used in this study is driven by the mouse *Prrx1* enhancer, but is easily modified to accommodate new tissue-specific regulatory elements. To facilitate uptake by other labs, we created a floxed INTACT cassette downstream of the ubiquitous *CAGGs* promoter (Fig. S3), making the INTACT*-olotl* workflow compatible with new and existing Cre recombinase strains (Currie et al., 2016; Gerber et al., 2018).

Applying the INTACT*-olotl* system over the course of pharmacological TGFβ inhibition allowed robust profiling of connective tissue states even from modest numbers of sequenced nuclei per timepoint. TGFβ pathway inhibition halted blastema formation, consistent with previous studies (Lévesque et al., 2007), but this block was temporary and removal of TGFβ inhibitor resulted in a resumption of blastema outgrowth. Transcriptionally, we observed large-scale decreases in ECM structural components and enzymes that are critical for remodeling of the wound site for eventual blastema formation. Notably, pro-regenerative components *Fn1* and *Tnc* (Ohashi et al., 2023; Rao et al., 2009) were downregulated, suggesting that the site of amputation is stalled at an early wound healing stage. TGFβ-inhibited CT-clusters highly express *Fap* (fibroblast activation protein), which is correlated with a pro-fibrotic/pro-inflammatory wound healing response (Fitzgerald and Weiner, 2020). This is further supported by the heightened expression of genes associated with interferon-gamma, interleukin, and tumor necrosis factor signaling. Together, this data suggests that TGFβ signaling is critical for resolving the wound site inflammation, and that this is necessary to progress through subsequent steps of the regenerative process. These snRNA-seq data suggest that TGFβ inhibition stalls connective tissue cells in a pro-fibrotic and pro-inflammatory environment reminiscent of mammalian wound healing (Tower et al., 2022). Overall, the INTACT-*olotl* system will provide an accessible and adaptable platform to further characterize cell type-specific transcriptomic and genomic mechanisms underlying complex tissue regeneration.

## Limitations

The INTACT*-olotl* workflow relies on labeling nuclei using a cell type-specific regulatory element. One caveat in doing so is that expression of the transgene can vary between targeted cells. Nuclei from cells expressing low levels of Sun1-sfGFP may therefore be captured inefficiently. Future iterations of INTACT-*olotl* will take advantage of floxed-INTACT transgenes that are driven from a ubiquitous CAGGs promoter (Fig. S3). Additionally, the INTACT system relies on labeling of nuclei and subsequent lysis of the cell membrane to liberate nuclei. Therefore, RNA present in the cytoplasm is lost during the process and is not available for downstream applications. Despite this, nuclear and whole-cell transcriptomes show high concordance in the expression of cell type and metabolic markers (Fig. S1) (Lake et al., 2017).

## Materials and Methods

### Animal husbandry, breeding, transgenesis, and drug treatments

Adult axolotls (*Ambystoma mexicanum*) with the d/d genetic background were maintained at Wake Forest University with standard husbandry practices as previously described (Khattak et al., 2013). The INTACT-*olotl* construct (*mPrrx1enh::Sun1sfGFP*, hereafter *mPrrx1enh::INTACT*) was created by subcloning a cassette containing a mouse N-truncated Sun1 nuclear membrane sequence tagged with sfGFP (Addgene Cat #160141, *pAAV-Ef1a-DIO-Sun1GFP-WPRE-pA*) (Lawler et al., 2020) into a plasmid containing the 2.4 Kb mouse *Prrx1* limb enhancer flanked by I-SceI and Tol2 integration sequences. Fertilized d/d embryos were injected with the INTACT-*olotl* plasmid as well as I-SceI meganuclease as previously described (Schuez and Sandoval-Guzmán, 2023). The resulting injected F0 animals were screened for the transgene fluorescence at the limb bud developmental stage 44/45 (Nye et al., 2003). F0 animals with both forelimb buds completely labeled were used for experiments as well as grown up to propagate F_1_ progeny. *mPrrx1enh::mCherry* axolotls used in FACS and subsequent sequencing experiments were obtained from an established transgenic line. All animals used for experiments were 3-6 cm in length from snout to cloaca. SB-431542 was purchased from Ambeed (CAS No.: 301836-41-9) and prepared according to Lévesque et al. (2007). A final concentration of 25µM SB-431542 was used to treat axolotls in 200 mL of Holtfreter’s housing water. Control animals received an equal volume of DMSO (equivalent to 0.1% of total volume) in Holtfreter’s solution which served as the vehicle only control. Solutions were refreshed daily and axolotls were housed under standard conditions. All experimental procedures were conducted in accordance with WFU IACUC (A23-118, A23-173, A25-208).

### Tissue collection for nuclei isolation

Prior to limb tissue collection, axolotls were anesthetized using a 0.007% benzocaine solution (Sigma-Aldrich). Subsequently, animals were transferred to a clean petri dish and moistened with anesthetic solution. The snout to cloaca length of each animal was measured and amputation was performed through the radius/ulna using a scalpel. Protruding bone was trimmed even to the amputation plane to promote subsequent regeneration. The amputated limb tissue was collected directly into cold hypotonic lysis buffer. The tissue was then immediately processed.

### Nuclei isolation and purification

Hypotonic lysis buffer solution B (Martin et al., 2023) with modifications was made fresh prior to each experiment and kept on ice throughout the process. 1mM DTT, 0.5mM spermidine, 0.1% Igepal, and 1x complete protease inhibitor EDTA-free (Roche) were added to PBS-based hypotonic buffer (Supplemental Material). RNase inhibitor was added at a final concentration of 1 U/µL to the lysis buffer and at a concentration of 0.5 U/µL in wash/resuspension buffer prior to use. Limb tissue was dissected into smaller pieces and placed into protein LoBind microcentrifuge tubes (Eppendorf) containing 1.5mL lysis buffer and one 5mm stainless steel bead. Homogenization was performed in a cooled 1600 MiniG homogenizer at 1400 rpm for 45-90 seconds depending on the size of limb tissue used. Samples were then incubated on ice for five minutes for lysis followed by two sequential passes through a 40µm cell strainer. Filtered suspension was centrifuged at 450rcf for five minutes at 4°C in a swinging bucket centrifuge to pellet nuclei. Supernatant was carefully removed, and the nuclei pellet was thoroughly resuspended in 500µL wash/resuspension buffer (Supplemental material). Sub-samples of resuspended nuclei were stained with 0.4% trypan blue to manually assess yield and overall morphology (i.e., intact nuclear membrane, lack of blebbing, absence of debris & aggregation).

Purification was performed using Chromotek GFP-Trap® M-270 magnetic particles. 5 µL of GFP-Trap beads were pre-cleared in low-RNase 1% BSA (Gibco) for 45 minutes at 4℃, rotating end over end. Beads were then washed three times and resuspended in wash/resuspension buffer. The Speed-IP protocol combined resuspended nuclei and GFP-Trap beads into a singular protein LoBind tube (Eppendorf) in a volume of 1.3 mL. The Purity-IP protocol distributed nuclei into three LoBind tubes with resuspended beads in equal final volume of 1.3 mL. Nuclei were then resuspended with a p1000 and rotated end over end for 30 minutes at 4℃ for the immunoprecipitation step. Tubes were then placed on a magnetic rack (New England Biolabs) and supernatant containing unbound nuclei was removed. The Speed-IP protocol used two washes, while the Purity-IP protocol used three total washes. Bead-bound *mPrrx1enh::INTACT^+^* nuclei were then resuspended in wash/resuspension buffer and quantified using 0.4% trypan blue staining. Subsequently, nuclei were directly used as input for snRNA-seq library prep using the Parse Biosciences WT mini v3 nuclei fixation kit.

### Tissue dissociation and FACS of *mPrrx1enh::mCherry* cells

The procedure for tissue collection, dissociation, and FACS was adapted from (Kawaguchi et al., 2024; Lin et al., 2021). Homeostatic and regenerating limb tissue from *mPrrx1enh::mCherry*-positive animals were collected under a stereoscope in a hood and washed twice in 1x Amphibian PBS (APBS) at room temperature. Subsequently, samples were dipped in 70% ethanol for 10 seconds and immediately washed twice in 1x APBS at room temperature. The tissue was dissected into pieces less than 0.5mm and immediately transferred to the enzymatic digestion solution (0.26WU/mL Liberase TM in 1x APBS, Roche). Enzymatic digestion was performed on a rotating wheel at room temperature for 45 minutes, followed by mechanical dissociation by pipetting cells up and down 10-15 times using a P1000 tip. Enzymatic digestion was stopped by adding an equal volume of high serum culture medium (10%FBS/AMEM: F12/DMEM, 10% FBS, 1% Anti-Anti, 1% ITS, 1% Glutamax, ddH_2_O). The resulting cell suspension was filtered sequentially through a 70µm and 30µm cell strainer. Dissociated cells were then pelleted by centrifugation at 300rcf for 5 minutes at room temperature. Isolated cells were washed twice with 1x APBS and then resuspended in 10% FBS/AMEM on ice. Fluorescence activated cell sorting (FACS) was performed using a Beckman-Coulter Astrios EQ cell sorter at the Wake Forest School of Medicine Flow Cytometry Shared Resource. *mCherry* positive cells were sorted using a 561 nm laser line and an 85µm nozzle. *mCherry^+^*cells were directly sorted into NucleoZOL solution (Macherey-Nagel, 740404.200) for subsequent bulk RNA isolation and purification.

### Nuclei isolation quality and positive nuclei quantification

To quantify the efficiency of the various nuclei isolation processes, we measured the area occupied by debris and nuclear aggregates in the field of a disposable hemocytometer (Corning). 10 µL of suspension from each condition was quantified in technical triplicate and repeated for three independent nuclei isolations. Images were acquired on a Zeiss Axio Zoom V16 using the brightfield filter with a total magnification of 112x. To visualize and quantify *mPrrx1enh::INTACT^+^* nuclei, nuclei were stained with DAPI (Thermo Fisher) and 5µL was placed on a #1.5 square coverslip. Nuclei were then imaged on a Zeiss LSM 880 inverted microscope using 405 and 488 nm laser lines. Positive nuclei were identified as having a distinct halo around the nucleus corresponding to the sfGFP inner nuclear membrane labeling. Nuclei for each condition were quantified using 3-5 technical replicates, where greater than 75 nuclei were counted each replicate. This process was repeated for at least three separate nuclei isolations and immunoprecipitations performed on different days.

### Image acquisition and histological tissue processing

Representative low magnification *in vivo* images showing expression of the INTACT cassette during limb development and homeostasis were captured using a Zeiss Axio Zoom V16 microscope. High magnification images of the limb bud, isolated nuclei, and distal blastema were captured on a Zeiss LSM 880 confocal microscope with the 20x or 63xw objective using 488nm laser excitation and transmitted light or DAPI to visualize overall nuclear morphology. Blastema tissue was harvested and fixed overnight in 4% PFA at 4℃, followed by equilibration in 30% sucrose/1xPBS. The tissue was then embedded in OCT (Thermo Fisher) and flash frozen on dry ice. Embedded tissue was sectioned at 14 µm and sections were collected on Colorfrost plus slides (Fisherbrand). Slides were then baked at 60℃ for 15 minutes to promote tissue adherence and remove excess OCT. Tissue sections were then washed with 1x PBS for 5 minutes at room temperature for three total washes. Sections were then briefly permeabilized in 1x PBST (0.1% Tween-20) for 15 minutes at room temperature followed by DAPI (Thermo Fisher) staining for 10 minutes. Slides were then washed once with 1x PBS and mounted using ProLong™ Glass Antifade Mountant (Invitrogen). Confocal images represent maximum intensity projections generated in ImageJ (20x developing limb bud substack = 15 µm, 63xw isolated nuclei substack = 9 µm, 20x blastema substack = 24 µm). Nuclei were spread on coverslips and if necessary immobilized with 1% agarose pads to prevent ambient movement and to ensure nuclei were all in a similar z-plane.

### RNA collection and bulk RNA sequencing

Limb tissue was collected under a stereoscope in an environment treated with RNase-Zap (Thermo, AM9780) followed by sanitization with 70% ethanol. Bulk limb tissue was subsequently placed in 500 µL of Nucleozol (Macherey-Nagel, 740404.200) solution with stainless steel homogenization beads on ice. Homogenization was carried out at 1400 rpm for five minutes at 4℃ using a 1600 MiniG instrument. Samples were stored at -80℃ or immediately processed. Nuclei were pelleted and resuspended in 500 µL Nucleozol solution. Total RNA was then isolated from samples using the Nucleospin RNA kit for NucleoZOL according to the manufacturer’s instructions (Macherey-Nagel, REF 740406.50). RNA was eluted in RNase-free H_2_O, quantified on a nanodrop spectrophotometer, and aliquoted and stored at -80℃.

Purified RNA was sent to Azenta Life Sciences for Illumina library preparation and sequencing at a depth of 50 million reads per sample. Reads were trimmed and aligned to the axolotl genome (UKY_AmexF1_1,2024, NCBI accession GCA_040938575.1) using Bowtie2. A count matrix was generated from BAM files using FeatureCounts. Counts were normalized to counts per million (CPM) and genes with CPM ≥ 1 were considered detected. Differentially expressed genes between whole tissue RNA and nuclear RNA were identified using DESeq2. Expression similarity between whole RNA and nuclear RNA samples was quantified by generating Spearman and Pearson correlations on a log2(CPM+1) scale (Fig. S1). Shared and unique genes between the two conditions were visualized on a Venn diagram (Fig. S1). GO analysis was then performed on the data using a custom axolotl curated GO term set identifying processes enriched in a given sample (FDR < 0.05) (Fig. S1).

### snRNA-seq experimental design and library preparation

Four F_1_ *mPrrx1enh::INTACT* hemizygous axolotls that were 4 cm in size (snout to cloaca) were distally amputated through the radius/ulna for each condition/timepoint. Axolotls were treated for eight consecutive days from the time of amputation with 25µM SB-431542 (TGFβ inhibitor) or equal volume of DMSO, with 200 mL solutions refreshed daily. Samples were collected at 5 and 10 days post amputation as well as a control uninjured sample from the zeugopod. Representative brightfield limb images were collected on a Zeiss Axio Zoom V16 using 25x total magnification. Regenerating tissue was collected from approximately 1mm behind the amputation plane to the distal tip. Tissue was split into two separate protein LoBind tubes with a singular 5mm stainless steel bead in each and were subjected to nuclei precipitation described above and then immediately fixed using the Parse Biosciences nuclei fixation kit according to manufacturer’s instructions. Nuclei were frozen in a Mr. Frosty isopropanol container (Thermo Fisher) overnight and stored at -80℃ until library preparation. Barcoded sublibraries (5000 nuclei each) were prepared to profile an equal number of nuclei per sample (Parse Biosciences Evercode WT v3 mini kit). Resulting libraries were quantified via Qubit and tapestation HS d5000 before sequencing. Sublibraries were sequenced by Novogene at a depth of 30,000 reads per nucleus (150 bp, paired end reads) using the Illumina NovaSeq X Plus instrument.

### snRNA-seq analysis

The Parse Biosciences Trailmaker pipeline was first used to generate a star index to an axolotl genome (UKY_AmexF1_1, NCBI accession GCA_040938575.1) with addition of a custom *mPrrx1enh::INTACT* sequence to map transgene mRNA expression. Raw FASTQ reads were then mapped to the axolotl star index using the Parse Biosciences pipeline (https://app.trailmaker.parsebiosciences.com/; Parse Biosciences Pipeline v1.7.3). Reads were then demultiplexed using Parse WT unique barcodes and aligned, trimmed reads were used to produce a gene expression count matrix. Within the Trailmaker insight module, the unfiltered data was processed by removing empty barcodes from the data using the trailmaker default threshold (Automatic, FDR = 0.01). Background was removed by manually selecting for presumptive nuclei with greater than 1000 transcripts per cell (Bin step = 200). The percentage of mitochondrial reads was quantified outside of the trailmaker pipeline using the R package Seurat. The average percentage of reads mapping back to mitochondrial encoded genes was below 0.1% for each sample, indicating that our nuclei isolation is largely devoid of cytoplasmic contamination, therefore nuclei were not filtered based on this criteria (Fig. S2). The Trailmaker pipeline then removed outliers from the genes vs. transcripts plot by fitting the data to a spline regression model (Automatic, Spline, Prediction interval = 0.999, p-value = 0.001). Doublets were then filtered out by using the scDblFinder method, using a probability threshold of 0.30 (Bin step = 0.0200). High quality nuclei passing these filters were then processed, normalized, and integrated using the Harmony R package (HVGs = 2000, lognorm, PCs = 20). On average we were able to identify 4,000-4,200 transcripts and 3,300-3,500 genes per nucleus across all timepoints and treatments (Fig. S2). Clusters were identified using the Leiden method and the data was visualized using a Uniform Manifold Approximation and Projection (UMAP) embedding (Min. distance = 0.9, cosine, Leiden, resolution = 0.3). Cluster specific marker genes were identified by analyzing the processed seurat object using the CyteType package Wilcoxon rank-sum test and LLM analysis (Ahuja et al., 2025). CyteType marker gene annotations were manually checked against previous marker gene sets and cell cluster predictions. Feature plots for the overall UMAP, gene set expression, and treatment groups were created in R using Seurat from the Parse Trailmaker exported filtered Seurat object. Seurat dot plots were created by subsetting average expression in CT-clusters only. Gene Ontology plots were generated using genes differentially expressed between DMSO and TGFβ-inhibited CT-clusters at 5 DPA. Genes with an adjusted p-value of 0.05 or less were used to calculate a gene ratio. GO terms were curated from NCBI RefSeq annotations (UKY_AmexF1_1, NCBI accession GCA_040938575.1) and significantly enriched terms were plotted based on having an adjusted p-value of less than or equal to 0.05. GO terms include biological processes, molecular functions, and cellular components. Implemented R code and additional materials used in this manuscript are available at the following GitHub repository: (https://github.com/currie-wfu/INTACT-olotl).

## Acknowledgements

We would like to thank members of the Currie lab for critical reading and feedback during execution of experiments and manuscript preparation. Thanks to Nick Leigh, Roger Deal, Ke Reid, Gloria Muday, and Nicholas Kortesis for feedback. This work was supported by a grant from the Eunice Kennedy Shriver National Institute of Child Health and Human Development NICHD 1R21HD119496-01 and Wake Forest University institutional startup funding to JDC. JEB and TC were partially supported through research assistantship funding from the Wake Forest Center for Molecular Signaling. We would like to thank the Wake Forest Biology Microscopic Imaging Core RRID:SCR_021975 for access and assistance with confocal and fluorescent stereoscope imaging, and the Wake Forest School of Medicine Cancer Genomics and Flow Cytometry Shared Resources.

