## Supplemental Figures for "*INTACT-olotl*: A simplified system for rapid cell type-specific profiling during axolotl limb regeneration"

Figure S1. **Comparison of bulk RNA sequencing results from total RNA vs. RNA from isolated nuclei.** (**A**) Correlation of gene expression from 14,503 genes shared between total RNA and nuclei RNA (Spearman's coefficient = 0.734, Pearson's coefficient = 0.749). Genes included in analysis had greater than 1 CPM. (**B**) Venn diagram visualization of genes shared between total RNA and nuclei RNA. Total RNA and nuclei RNA had 2,361 and 785 unique genes detected, respectively. (**C-D)** GO analysis of genes detected only in total RNA and nuclei RNA. The most represented GO term in the total RNA sample corresponds to the plasma membrane (underlined). The top two represented GO terms in the nuclei RNA sample correspond to RNA polymerase DNA binding (underlined).

Figure S2. **Sn-RNAseq quality control and validation of TGFβ pathway inhibition.** (**A**) % of total reads mapping back to mitochondrial encoded genes across all samples. (**B**) Number of transcripts detected represented on a log10 scale across all samples. (**C**) Number of genes identified represented on a log10 scale across all samples. (**D**) UMAP visualization of *Sun1-sfGFP* mRNA (transgene) expression across all cell clusters. UMAP represents all samples and timepoints. Note transgene expression in the bottom left clusters corresponding to CT cell types. (**E**) Dot plot depicting changes in the average expression of TGFβ pathway components across all timepoints (0, 5, 10 DPA) and conditions (DMSO, TGFβ Inh.) within the CT cluster's. CT clusters' under TGFβ inhibition at 5 DPA significantly downregulated TGFβ pathway components relative to the 5 DPA DMSO control. Expression of TGFβ pathway components is restored once samples have been washed out of the inhibitor for two days.

Figure S3. **Floxed INTACT cassette driven by ubiquitous CAGGs promoter.** (**A**) Schematic of transgene containing the **γ**-Crystallin promoter driving expression of EYFP within the eye before recombination, allowing for efficient screening of positive animals. Upon cell-type specific Cre recombination, the INTACT cassette is driven by the ubiquitous CAGGs promoter. (**B-D**) Visualization of **γ**-Crystallin promoter driven EYFP expression in the eye of a non-recombined axolotl. There is no leaky expression of the INTACT cassette within the limb of positive axolotls prior to Cre recombination.

**Other Supplementary Information:**

Supplemental Material 1 - **Nuclei isolation and immunoprecipitation solutions for Speed/Purity-IP**

### Figure S1

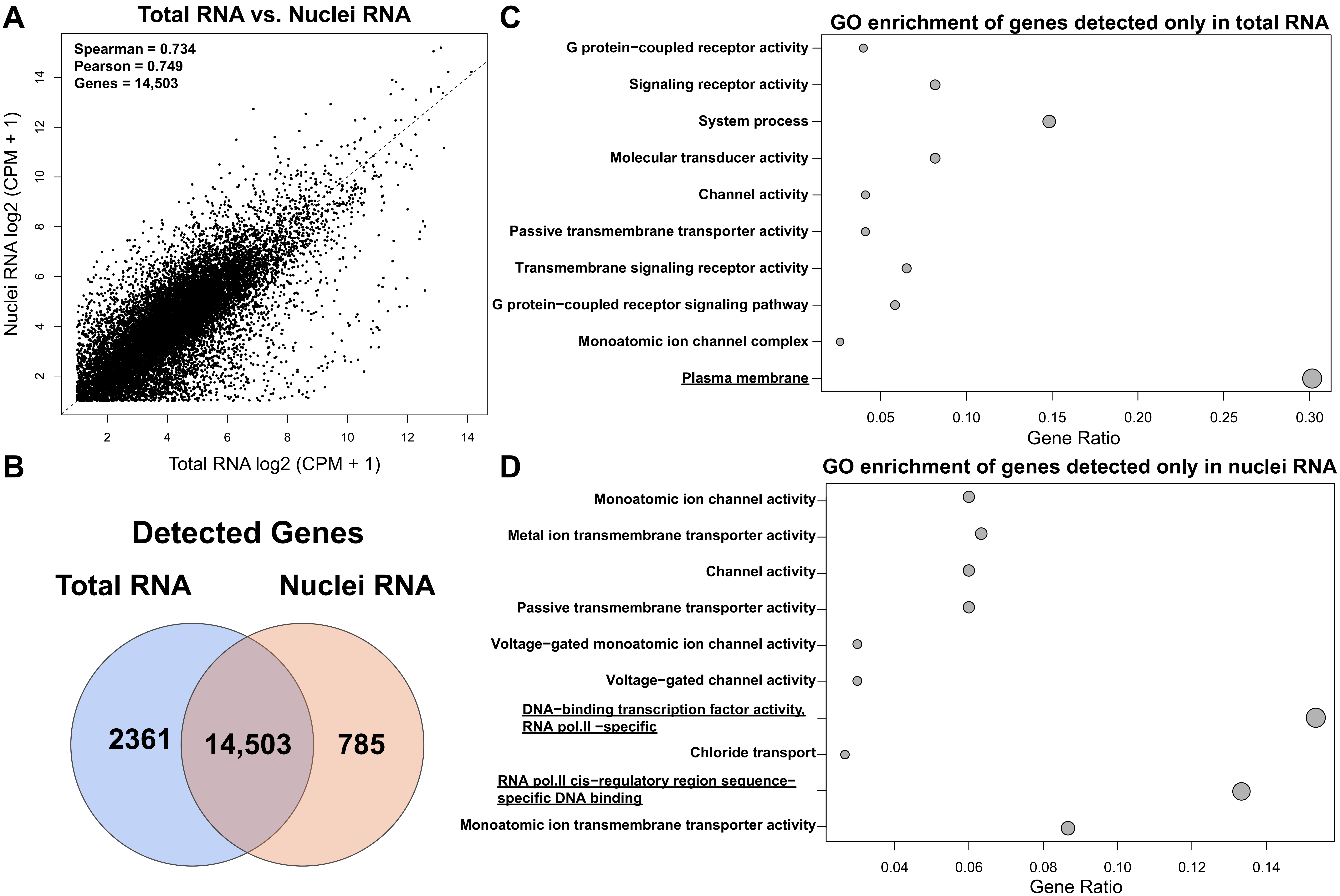

A
Total RNA vs. Nuclei RNA
Spearman = 0.734
Pearson = 0.749
Genes = 14,503
Nuclei RNA log2 (CPM + 1)
Total RNA log2 (CPM + 1)
2
4
6
8
10
12
14
B
Detected Genes
Total RNA
Nuclei RNA
2361
14,503
785
C
GO enrichment of genes detected only in total RNA
G protein−coupled receptor activity
Signaling receptor activity
System process
Molecular transducer activity
Channel activity
Passive transmembrane transporter activity
Transmembrane signaling receptor activity
G protein−coupled receptor signaling pathway
Monoatomic ion channel complex
Plasma membrane
0.05
0.10
0.15
0.20
0.25
0.30
Gene Ratio
D
GO enrichment of genes detected only in nuclei RNA
Monoatomic ion channel activity
Metal ion transmembrane transporter activity
Channel activity
Passive transmembrane transporter activity
Voltage−gated monoatomic ion channel activity
Voltage−gated channel activity
DNA−binding transcription factor activity, RNA pol.II −specific
Chloride transport
RNA pol.II cis−regulatory region sequence−specific DNA binding
Monoatomic ion transmembrane transporter activity
0.04
0.06
0.08
0.10
0.12
0.14
Gene Ratio

Figure S2

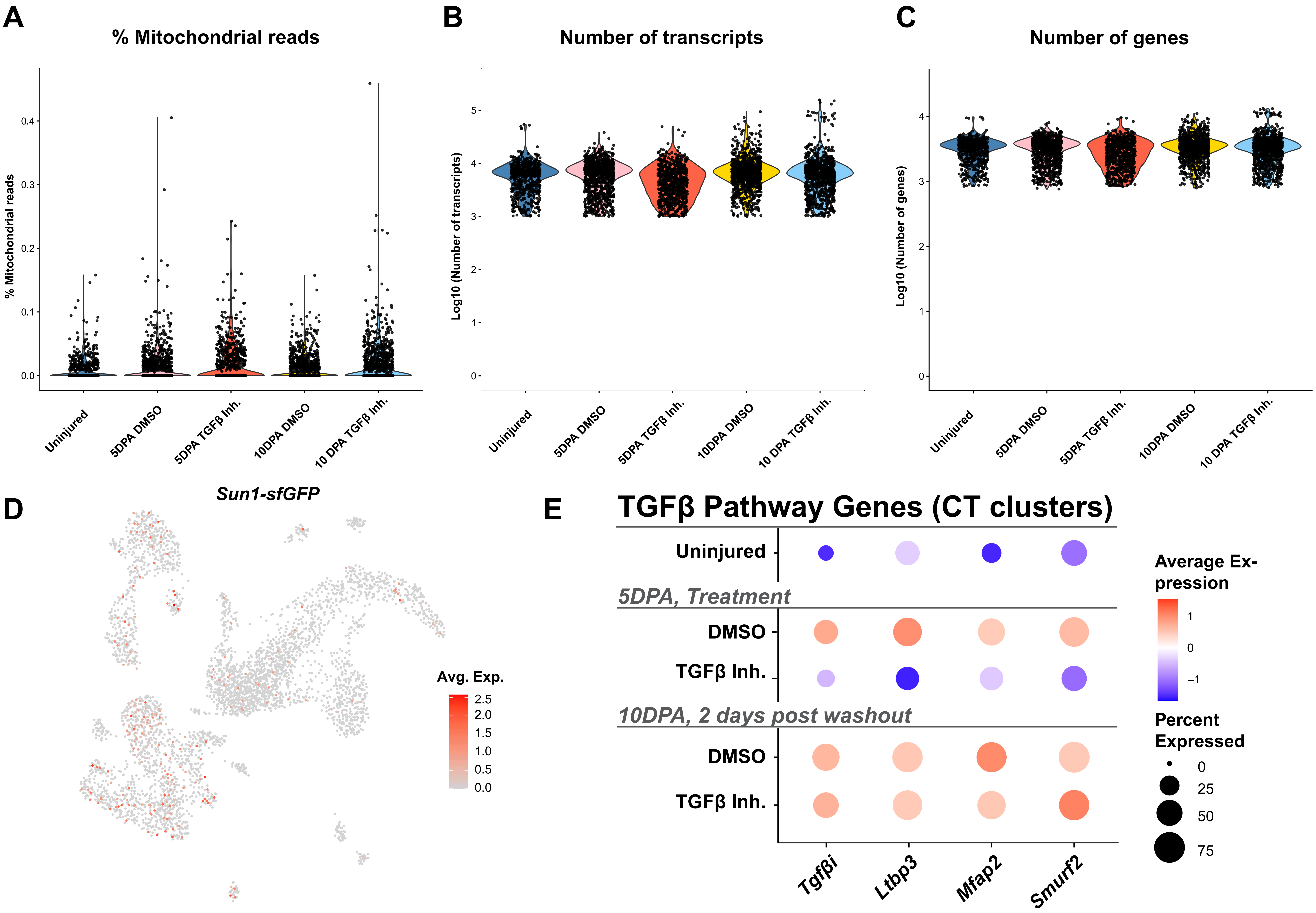
A
% Mitochondrial reads
% Mitochondrial reads
0.0
0.1
0.2
0.3
0.4
Uninjured
5DPA DMSO
5DPA TGFβ Inh.
10DPA DMSO
10 DPA TGFβ Inh.
B
Number of transcripts
Log10 (Number of transcripts)
0
1
2
3
4
5
Uninjured
5DPA DMSO
5DPA TGFβ Inh.
10DPA DMSO
10 DPA TGFβ Inh.
C
Number of genes
Log10 (Number of genes)
0
1
2
3
4
Uninjured
5DPA DMSO
5DPA TGFβ Inh.
10DPA DMSO
10 DPA TGFβ Inh.
D
Sun1-sfGFP
Avg. Exp.
2.5
2.0
1.5
1.0
0.5
0.0
E
TGFβ Pathway Genes (CT clusters)
Uninjured
5DPA, Treatment
DMSO
TGFβ Inh.
10DPA, 2 days post washout
DMSO
TGFβ Inh.
Tgfβi
Ltbp3
Mfap2
Smurf2
Average Expression
1
0
−1
Percent Expressed
0
25
50
75

### Figure S3

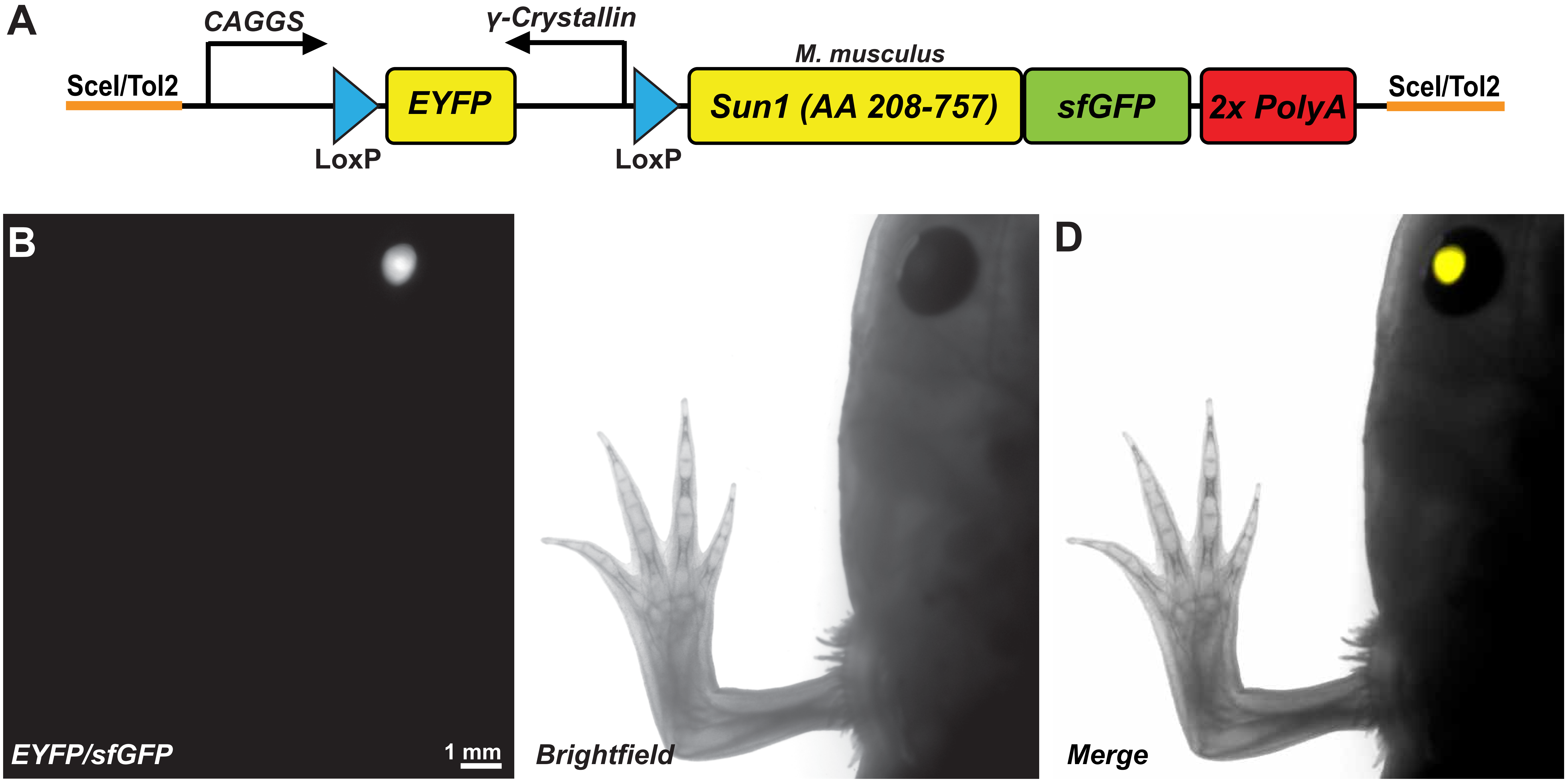
A
CAGGS
γ-Crystallin
M. musculus
SceI/Tol2
EYFP
Sun1 (AA 208-757)
sfGFP
2x PolyA
SceI/Tol2
LoxP
LoxP
B
EYFP/sfGFP
1 mm
Brightfield
D
Merge
