## Supplemental Material 1 for "*INTACT-olotl*: A simplified system for rapid cell type-specific profiling during axolotl limb regeneration"

**Nuclei isolation and immunoprecipitation solutions**

**10x PBS-hypotonic stock:**

| **Reagent** | **Amount (g)** | **Concentration (mM)** |
| --- | --- | --- |
| Na_2_HPO_4_ (dibasic) | 5.45 | 76 |
| NaH_2_PO_4_ . H_2_O | 3.1 | 44 |
| KH_2_PO_4_ | 1.2 | 17 |
| KCl | 1 | 26 |
| NaCl | 3 | 102 |

-Fill to 500 mL with DEPC-treated water, sterile filter. Store at room temperature for up to 6 months.

**Lysis base buffer solution:**

- 5 mL 10x PBS-hypotonic stock
- 5.7 g sucrose (333 mM)
- 75 μL 2M MgCl_2_ (3 mM)
- Fill to 50 mL with DEPC-treated water

Sterile filter, store at 4 °C for up to a month

**1x PBS-hypotonic lysis buffer:**

| **Solution** | **Volume (μL)** | **Concentration** |
| --- | --- | --- |
| Lysis base buffer solution | 3,557.5 | 1x |
| 7.5% BSA | 670 | 1% |
| 10% Igepal | 50 | 0.1% |
| 1M DTT | 5 | 1 mM |
| 1M Spermidine | 2.5 | 0.5 mM |
| 7x cOmplete mini EDTA-free protease inhibitor (Roche) | 715 | 1x |

-1x PBS-hypotonic lysis buffer was made fresh right before use and stored on ice.

**Modified SPBSTM stock (Wash and resuspension buffer):**

- 28.5 g sucrose (333 mM)
- 375 μL 2M MgCl_2_ (3 mM)
- Fill up to 250 mL with 1x PBS (using DEPC-treated water)

**Modified SPBSTM working solution:**

| **Solution** | **Volume (μL)** | **Concentration** |
| --- | --- | --- |
| SPBSTM Stock | 4,305 | 1x |
| 7.5 % BSA | 670 | 1% |
| 10% Igepal | 25 | 0.05% |

*RNase inhibitor was added to 1x hypotonic lysis buffer at a final concentration of 1U/μL and 0.5 U/μL in SPBSTM wash/resuspension buffer prior to use*
